# How cortico-basal ganglia-thalamic subnetworks can shift decision policies to maximize reward rate

**DOI:** 10.1101/2024.05.21.595174

**Authors:** Jyotika Bahuguna, Timothy Verstynen, Jonathan E. Rubin

## Abstract

All mammals exhibit flexible decision policies that depend, at least in part, on the cortico-basal ganglia-thalamic (CBGT) pathways. Yet understanding how the complex connectivity, dynamics, and plasticity of CBGT circuits translate into experience-dependent shifts of decision policies represents a longstanding challenge in neuroscience. Here we present the results of a computational approach to address this problem. Specifically, we simulated decisions driven by CBGT circuits under baseline, unrewarded conditions using a spiking neural network, and fit an evidence accumulation model to the resulting behavior. Using canonical correlation analysis, we then replicated the identification of three control ensembles (*responsiveness*, *pliancy* and *choice*) within CBGT circuits, with each of these subnetworks mapping to a specific configuration of the evidence accumulation process. We subsequently simulated learning in a simple two-choice task with one optimal (i.e., rewarded) target and found that feedback-driven dopaminergic plasticity on cortico-striatal synapses effectively manages the speed-accuracy tradeoff so as to increase reward rate over time. The learning-related changes in the decision policy can be decomposed in terms of the contributions of each control ensemble, whose influence is driven by sequential reward prediction errors on individual trials. Our results provide a clear and simple mechanism for how dopaminergic plasticity shifts subnetworks within CBGT circuits so as to maximize reward rate by strategically modulating how evidence is used to drive decisions.

**Author summary:** The task of selecting an action among multiple options can be framed as a process of accumulating streams of evidence, both internal and external, up to a decision threshold. A decision policy can be defined by the unique configuration of factors, such as accumulation rate and threshold height, that determine the dynamics of the evidence accumulation process. In mammals, this process is thought to be regulated by low dimensional subnetworks, called control ensembles, within the cortico-basal ganglia-thalamic (CBGT) pathways. These control ensembles effectively act by tuning specific aspects of evidence accumulation during decision making. Here we use simulations and computational analysis to show that synaptic plasticity at the cortico-striatal synapses, mediated by choice-related reward signals, adjusts CBGT control ensemble activity in a way that improves accuracy and reduces decision time to maximize the increase of reward rate during learning.

## Introduction

A characteristic of nearly all mammals is the ability to quickly and flexibly shift how currently available evidence is used to drive actions based on past experiences (1). For example, feedback may be used to shift between making exploratory decisions, where low value actions are sampled to gain information, and exploitative decisions, where high value actions are taken to maximize immediate rewards (2; 3; 4). Orthogonal to this exploration-exploitation dimension is a complementary choice about decision speed: actions can be made quickly or slowly depending on immediate goals and confidence level (5). These shifts between fast or slow and exploratory or exploitative decision policies can be interpreted as different states of an underlying evidence accumulation process (6; 7), often captured by mathematical models such as the drift diffusion model (DDM; (8; 9; 10; 11; 12)). From this perspective, the values of DDM parameters, such as the drift rate (*v*; the rate of evidence accumulation during a single decision) and boundary height (*a*; the amount of evidence needed to trigger a decision) can be tuned to capture a particular decision policy. Thanks to this mapping, specific (*a, v*) pairs effectively correspond to positions on a manifold of possible decision policies that determine how both internal and external evidence combine to drive eventual actions (Figure 1, ”WHAT” panel). Although speed and accuracy are negatively correlated *a priori*, according to Fitt’s Law (13), the goal of learning is to converge to a position on this manifold of decision policies that manages to optimize both speed and accuracy for a given task context (14; 15; 16).

**Fig 1.**
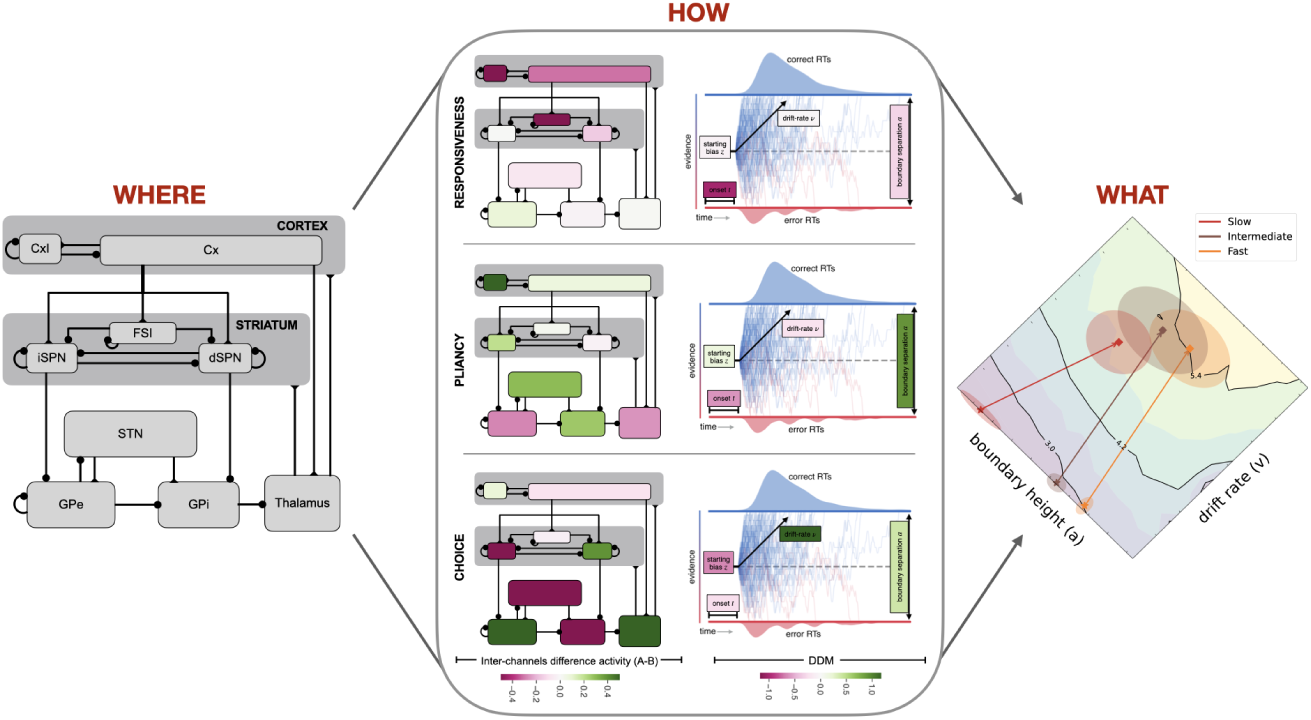
Decision-making deconstructed. Most voluntary decision policies depend on the CBGT circuits (WHERE; left panel). This can be described at the algorithmic level by a set of parameters in a process model (e.g., the DDM) that drives an evidence accumulation process. The goal of this process is to determine the effective reward rate of choices (WHAT; right panel contours), as well as other decision parameters. Control ensembles within CBGT circuits determine the relative configuration of decision policy parameters (HOW; middle panel) (30). What remains unclear, and we address in this work, is how learning modulates the balance between control ensembles in a way that shifts decision policies so as to maximize reward rate. Cx, cortical PT cells; CxI, inhibitory interneurons; FSI, fast spiking interneurons; d/iSPN, direct/indirect spiny projection neurons; STN, subthalamic nucleus; GPe, external globus pallidus; GPi, internal globus pallidus CBGT network were modeled across stages of learning using the DDM (see (29; 30**?** )).

This form of learning is managed, at least in part, by the cortico-basal ganglia-thalamic (CBGT) circuits, a distributed set of interconnected brain regions that is ideally situated to influence nearly every aspect of decision-making (17; 18; 19; 20; 21) (Fig. 1, ”WHERE” panel). The cannonical CBGT circuit includes a collection of interacting basal ganglia pathways that receive cortical inputs and compete for control of an output region (predominantly the internal globus pallidus, GPi, in primates or the substantia nigra pars reticulata, SNr, in rodents) that impacts thalamocortical or superior collicular activity to influence actions (22; 23; 24). The balance of this competition is thought to map to a configuration of the evidence accumulation process (7; 25; 26; 27; 28; 29; 30). Therefore, if behavioral flexibility reflects the *what* and CBGT circuits represent the *where* of flexible decision-making, then we are left with an open question of *how* : how do CBGT circuits control flexibility in decision policies during learning?

In prior work we showed how the computational logic of normative CBGT circuits can be expressed in terms of three low-dimensional subnetworks, called *control ensembles*. Each control ensemble tunes specific configurations of the evidence accumulation process, manifested as control over distinct dimensions of a decision policy (30). In theory, these control ensembles, dubbed *responsiveness*, *pliancy*, and *choice* (Fig 1, ”HOW” panel), provide candidate mechanisms for implementing shifts in decision policies during learning. Here we illustrate how a single plasticity mechanism acting at the cortical inputs to the basal ganglia can, through network interactions, leverage the control ensembles to steer behavior during learning. To this end, we simulated a biologically-constrained spiking CBGT model that learns to select one of two actions via dopamine-dependent plasticity, driven by reward prediction errors, at the cortico-striatal synapses. We then implemented an upwards mapping approach (31), in which the behavioral features (decision times and choices) produced by the simulated Finally, we used various analytical approaches to replicate the existence of the low-dimensional control ensembles prior to learning and quantify how their influence levels change over the course of training. Our results show that value-based learning leads to a specific tuning of CBGT control ensembles in a way that manages the speed-accuracy tradeoff so as to maximize reward rate across successive decisions.

## Results

### Feedback learning in CBGT networks maximizes reward rate

We consider a situation where an agent encounters a new environment for which it has no relevant prior experience or bias, so that the selection of all options is equally likely at first. In a simple two-choice bandit task, with one rewarded and one unrewarded option, this unbiased starting point would correspond to a 50% error rate. With learning it should be possible to make fewer errors over time, leading to increased rewards, but exactly how this is achieved in practice depends on the decision policy that the agent adopts. For example, if the agent prioritizes speed over all else in its action selection, then its error rate will likely remain high, leading to fewer rewards over time. Conversely, by making sufficiently slow decisions, the agent may be able to achieve an extremely low error rate, leading to greater likelihood of reward on individual trials, but if the response speed is too slow then the rate of reward return over time may take a significant hit. The overall reward rate achieved by the agent thus depends on both decision speed and accuracy. Intuitively this may be optimized for a fixed level of experience via some compromise between these two dimensions (14; 32).

To understand how optimized speed and accuracy emerge from CBGT circuits, we simulated 300 instances of a spiking computational model of CBGT pathways. For each instance, a parameter set was pseudo-randomly selected from preset intervals to keep average firing rates of all relevant cell types within known biological ranges (updated slightly from our past work (30); see Supporting Information Appendix, Supp. Fig. S1A). The networks performed a two-armed bandit task with deterministic reward feedback (i.e., the reward probability was 100% for the optimal choice and 0% for the suboptimal one). Using a deterministic reward task, as opposed to a task where rewards are delivered probabilistically, explicitly ties accuracy to reward return and also makes the optimal learning strategy a simple “win-stay/lose-switch” policy (33). Learning in the network was implemented with dopamine-dependent plasticity at the cortico-striatal synapses, where the magnitude of the phasic dopamine response following each decision was based on reward prediction error (for details see (34)). It should be noted that despite this being a deterministic task, there is ample variability in the network performance. Even after 15 trials of learning, the networks do not reach perfect performance, averaging ≈ 90% accuracy (Supp. Fig. S3).

Following a set of simulated trials in this task, we fit the reaction time (RT) and choice probabilities of each network, together reflecting the form of its decision policy, with a hierarchical version of the DDM (35; 36). The DDM provides an intuitive framework for mapping behavioral responses to an evidence-accumulation representation of the decision policy that can be described by only a few parameters (8). Although there are many possible variants of the DDM that we could use, including versions with collapsing bounds (37; 38; 39) or trial wise evolution of specific parameters (40), our task does not involve factors like urgency or non-stationarity of task states. Thus, we opted for the simplest version of the model for the sake of parsimony. The quality of the fits obtained are shown in Supp Fig. S2. After each predetermined step in learning (2, 4, 6, and 15 trials with plasticity on), we froze the network by turning off plasticity, simulated 300 trials to generate an RT distribution and choice probabilities from the current state of the network, and fit the DDM to these behavioral measures again. After these probe trials, learning was turned back on and the task progressed. This process allowed us to plot each network’s performance as a trajectory in the DDM parameter space.

Because of the mapping from network behavioral responses to DDM parameters, we will refer to the 2-dimensional plane of drift rates (*v*) and boundary heights (*a*) as a decision policy manifold. Figure 2 shows the average trajectories of three groups of networks on the (*v, a*) decision policy manifold. For each *v* and *a* we also estimated the average RT (Fig. 2A), accuracy (Fig. 2B) and reward rate (Fig. 2C; see also (15)). The three network groups shown in this figure represent a tertiary split of the full set of simulated networks into fast (short RT, orange), intermediate (medium RT, brown), and slow (long RT, red) groups, based on their initial RT values (Fig. S1B). We implemented this split to determine whether decision policy adjustments due to learning were influenced by initial biases in the networks. Despite their initial speed differences, all three network groups showed chance level performance before plasticity (Fig. S1C) and converged to similar regions of the (*v, a*) space with learning (Figure 2, shaded ellipses). A comparison of behavioral measures and DDM parameters before and after plasticity is presented in Figure S3.

**Fig 2.**
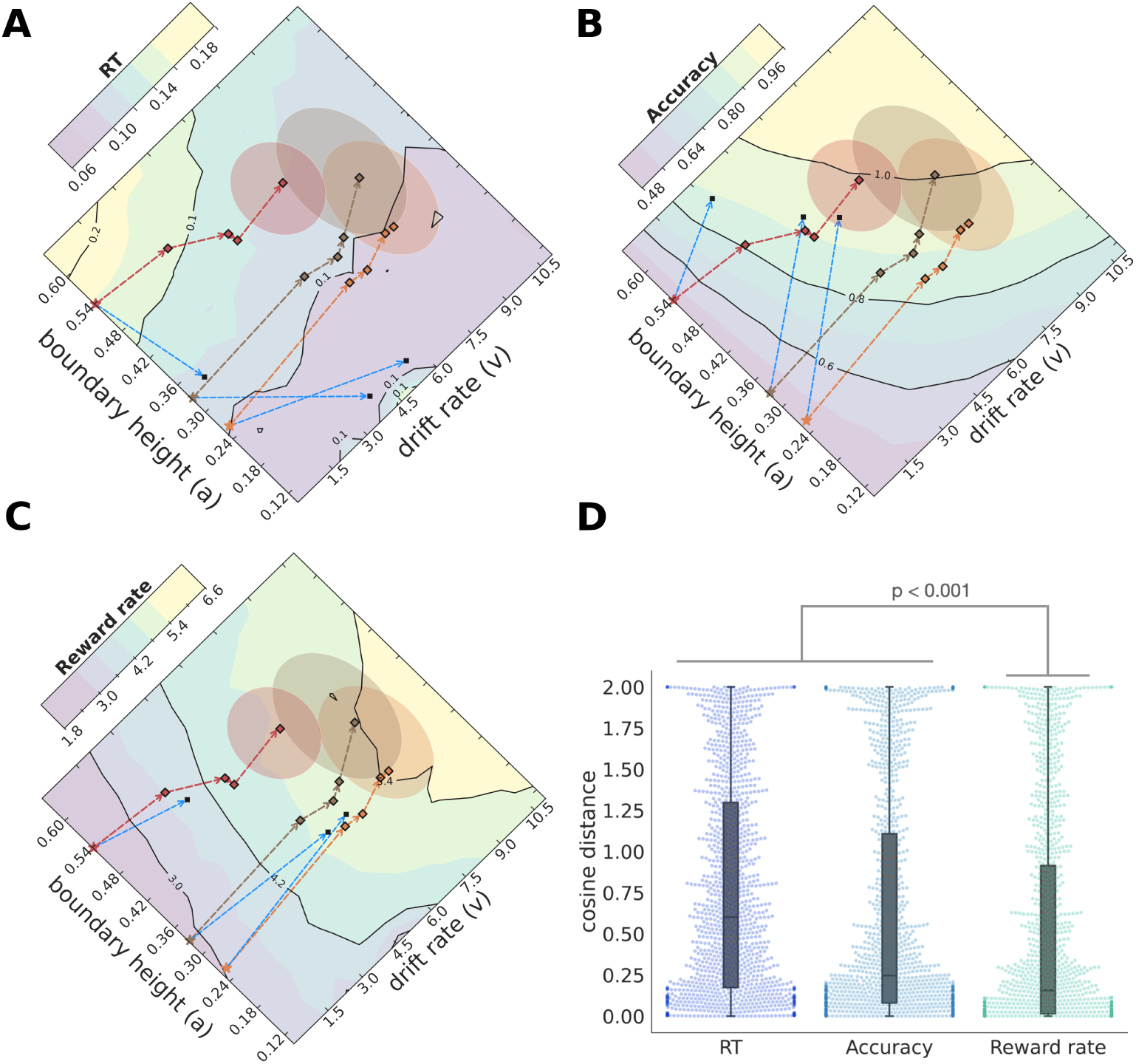
Dopamine-dependent cortico-striatal plasticity drives CBGT networks in the direction of reward rate maximization. (A) The evolution of RTs achieved by a DDM fit to CBGT network behavior, projected to (*v, a*)-space. The average starting position for the fast (orange), intermediate (brown) and slow (red) networks are shown as stars. The squares indicate the evolution of each network group over the plasticity stages, which converge after 15 trials (shaded elliptical regions). The yellow (purple) colors represent high (low) RTs. The network trajectories do not evolve in the direction that would be expected to minimize the RTs (e.g., optimal direction shown in blue from the initial position of all three speed groups). (B) The yellow (purple) colors represent high (low) accuracy. The networks evolve towards increasing expected accuracy but not in an optimal fashion (trajectories vs. blue arrows). (C) The yellow (purple) colors represent high (low) reward rate. The network evolution aligns closely with the direction that maximizes the reward rate (blue arrows). (D) The cosine distances calculated for every network at each plasticity stage for RT, accuracy and reward rate are were pooled together and shown as distributions.

These trajectories clearly demonstrate that our CBGT network can learn from simple dopaminergic feedback at the cortico-striatal synapses. But what exactly is the objective being maximized by the network? To address this question, we compared the change at each step of learning to the predicted direction that the network would take if it were maximizing one of the three possible behavioral objectives: speed, accuracy, or reward rate. Note that we can plot contours for each of these quantities (using RT as a gauge of speed) over the (*v, a*) domain. Although the mapping between (*v, a*) and either speed, accuracy or reward rate is not bijective, we will nonetheless refer to the (*v, a*) plane as the speed manifold, accuracy manifold, or reward rate manifold when it is shown along with the contours of the corresponding quantity. These predicted directions of objective maximization are illustrated as blue vectors in Figure 2A-C, reflecting steps from each initial point that are in the direction of the gradient of each objective (i.e., the direction of maximal change, which lies orthogonal to the contours, shown with the same length as the vector representing the actual network evolution at the first step of learning in each case). Analysis of the trajectories in Figure 2A reveals that while plasticity decreases RTs with learning, the angles of the learning trajectories do not align with the optimal directions for maximally reducing RT. Similarly, the network trajectories do not align with the vectors that would be expected if they were maximizing accuracy alone (Figure 2B). In contrast, the average trajectories along the reward rate manifold were closest to the gradient and hence to the optimal trajectories attainable for that manifold (Fig. 2C). Moreover, the rate of increase in reward rate was similar regardless of the network’s initial speed bias.

To quantify the alignment of observed network trajectories to the expected directions of maximal change, we calculated the cosine distance between the observed vector and the optimal vector, normalized to the observed vector’s length, at each learning step. While there is substantial variability across networks (Figure 2D), there was a consistent effect of objective type on the network fits (F[3597, 2]=47.2, p*<*0.0001). Fits to the reward rate trajectories were consistently better than to either speed (t(299)=13.59, p*<*0.0001) or accuracy (t(299)=8.35, p*<*0.0001) trajectories. This effect held regardless of a network’s initial speed (Figure S4). Thus, our biologically detailed model of the CBGT circuit can effectively learn to maximize reward rate by managing the speed-accuracy tradeoff during the evidence accumulation process via dopaminergic plasticity at the cortico-striatal synapses.

### Low-dimensional control ensembles that map to general decision policies

The CBGT network and DDM are, respectively, implementation-level and algorithmic-level descriptions of the evidence accumulation process that guides goal-directed behavior. We have previously shown that there is a low-dimensional, multivariate mapping between these two levels of analysis in the absence of learning (30). Here we set out to replicate this observation with the CBGT parameter sets used in the current study, with the aim of analyzing their contributions to the dopaminergic learning process. For this step, we considered two aspects of activity within each CBGT population: global activation across the two action representations (sum of the activity in that region, across both channels; Σ) and bias towards one action representation (difference in activity within each region, across the action channels; Δ). Using canonical correlation analysis (CCA), we captured the low-dimensional components that maximally correlate variation in CBGT activity with variation in DDM parameters. This analysis identified three such components (Fig. 3). We refer to these low-dimensional components as *control ensembles*.

**Fig 3.**
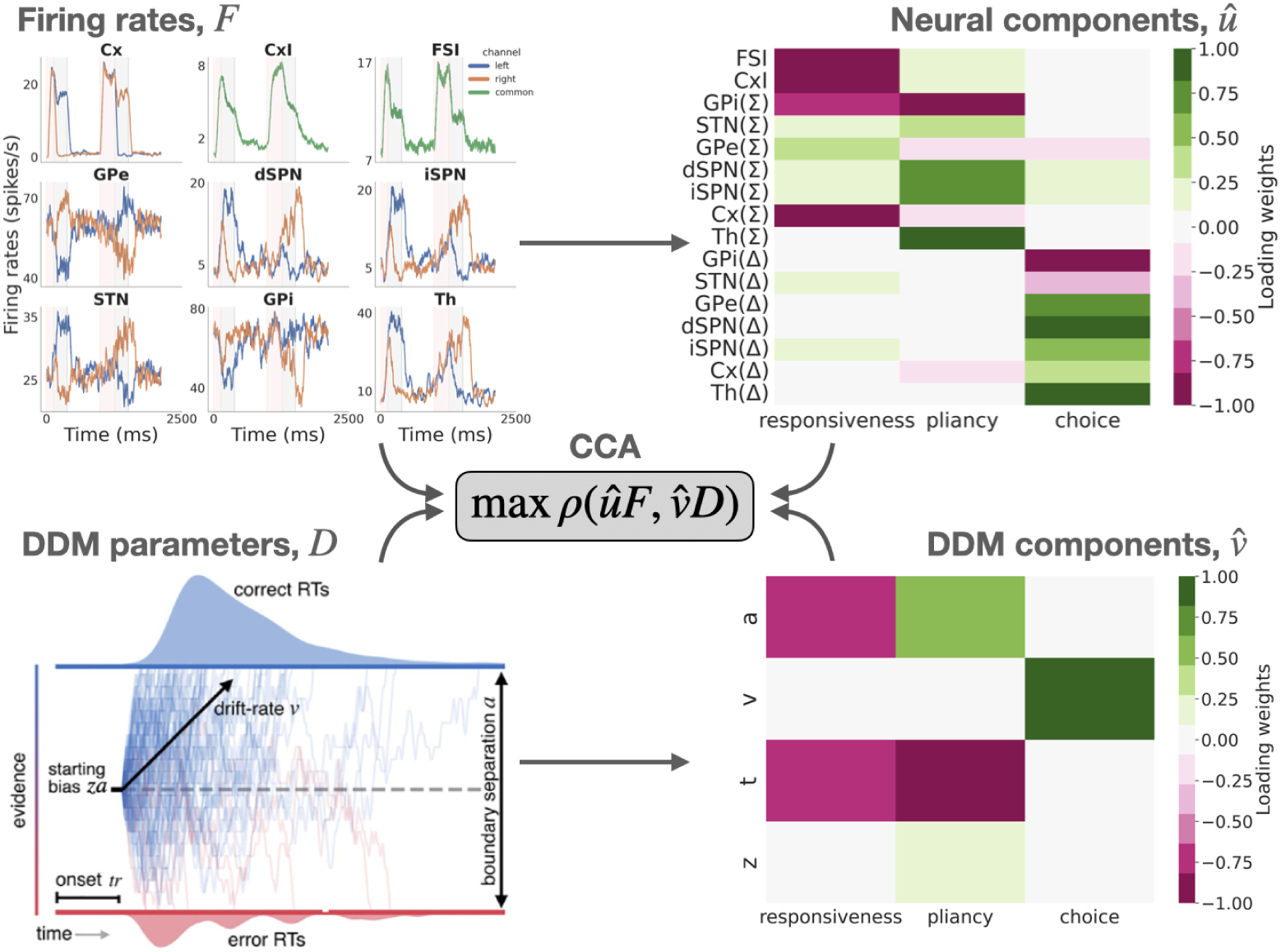
Canonical correlation analysis (CCA) identifies control ensembles (cf. (**30**)). Given matrices of average firing rates, *F* (both summed rates across channels, Σ, and between-channel differences, Δ), and fit DDM parameters, *D*, derived from a set of networks at baseline (left panels), CCA finds the low-dimensional projections, *u*^ for firing rates and *v*^ for DDM parameters (right panels), which maximize the correlation, *ρ*, between the projections *u*^*F* and *v*^*D* of *F* and *D*. Blue lines in the *F* plot show left channel activity, orange show right channel activity, and green shows populations that go across both channels.

The three control ensembles identified by our analysis nearly perfectly replicate our prior work (30), where they are described in more detail (see also Section *Upward mapping* ). Thus we kept the labels *responsiveness*, *pliancy*, and *choice* ensembles for the first, second, and third components, respectively. The recovered components are shown in both CBGT and DDM parameter spaces in Figure 3 (right panels). The responsiveness component describes the agent’s sensitivity to evidence, both in terms of the delay before the agent starts to accumulate evidence (*t*) and how significantly the presence of evidence contributes to achieving the decision threshold (*a*). The dominant features of CBGT activity that vary along the responsiveness control ensemble loadings are a global inhibitory signal, including fast-spiking interneuron (FSI) and overall internal globus pallidus (GPi(Σ)) activity, as well as overall excitatory and inhibitory cortical activity (Cx(Σ), CxI). Because the dominant CBGT and DDM loadings for the responsiveness control ensemble have the same sign (all negative), they imply that a *decrease* in the weighted activity of the loaded populations corresponds to an *decrease* in onset time, *t*, and *a* and, hence, to an *increase* in overall responsiveness.

The pliancy component refers to the level of evidence that must be accumulated before committing to a decision. As with responsiveness, pliancy loads mostly on *a* and *t*, but now with opposing signs for these two loadings, corresponding to the idea that even though an agent is attentive to evidence (small *t*), it requires a substantial accumulation of evidence to reach its threshold (large *a*). The CBGT activity features that characterize pliancy are the overall engagement of the BG input nodes (i.e., global dSPN and iSPN activity, with a smaller STN contribution), as well as total GPi and thalamic activity, with oppositely signed loadings to each other. For the pliancy component, a change in the activity consistent with the cell type loadings (e.g., increase in SPN activity) corresponds to a decrease in overall pliancy (e.g., increase in *a*).

Lastly, the choice component represents the intensity of the choice preference and is reflected largely in *v* and the neural correlates of competing choice representations in the CBGT (i.e., differences in activity across the two action channels within each BG region). A change in activity consistent with the cell type loadings (e.g., greater difference in dSPN activity between the two channels) corresponds to a stronger commitment towards the more rewarded option (i.e., larger *v*).

In summary, each CBGT control ensemble can be interpreted as specifying a coordinated collection of changes in CBGT neural activity levels that can, in theory, most effectively tune a set of decision policy parameters (captured by the DDM). Now that we have delineated the control ensembles embedded within the CBGT network (cf. (30)), we are ready to consider how dopamine-dependent plasticity regulates their influence in a way that collectively drives decision policies to maximally increase reward rate.

### Cortico-striatal plasticity drives control ensembles during learning

Our analysis of the CBGT network behavior (Figure 2) shows that dopamine signaling at the cortico-striatal synapses is enough to elicit changes in the evidence accumulation process that maximize reward rate. This observation suggests that there are emergent driver mechanisms, originating from cortico-striatal synaptic changes, that tune the control ensembles in a way that achieves this outcome. That is, if each control ensemble represents a knob to tune an aspect of the decision policy, then a driver mechanism selects a set of adjustments of the knobs that yields an overall decision policy selection. We next set out to identify these emergent drivers.

As a first step toward quantifying the modulation of CBGT activity after 15 learning trials, we calculated the principal components of the overall change in firing rates of all 300 networks. The first 5 of these components collectively explain more than 90% of the observed variance (Fig. S5A, thick blue line marked ”All”). The loading weights (Fig. 4A) show that the first and third components reflect the global activity of subsets of CBGT nuclei. The second, fourth and fifth components relate more strongly to the bias towards one option, with predominant loadings on differences in rates across action channels. Together, these components represent the collection of changes in firing rates that result from learning-related changes at the cortico-striatal synapses.

**Fig 4.**
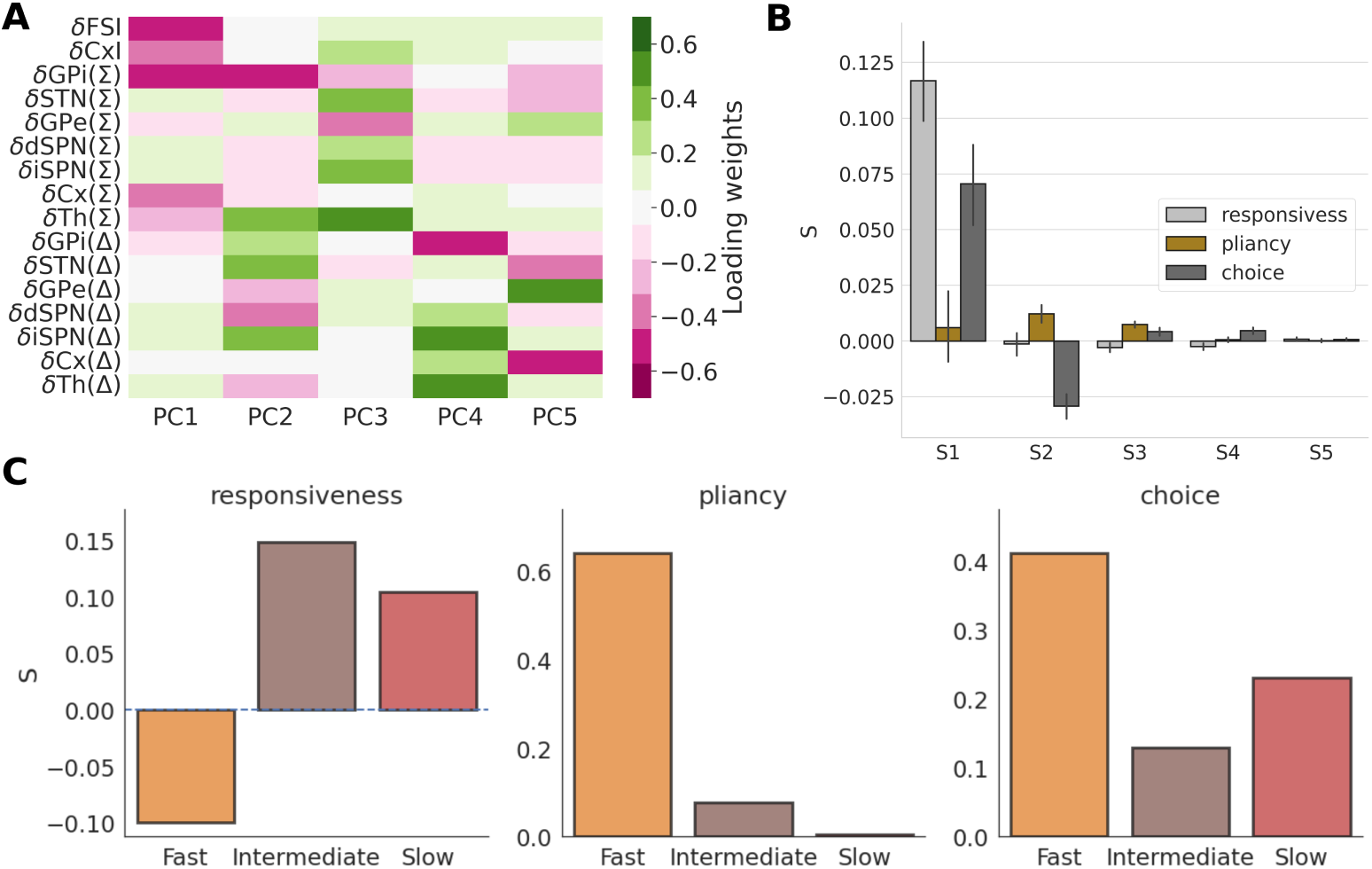
Plasticity-induced changes of control ensemble influence. (A) The loading weights of the first 5 PCs of firing rate changes from before to after plasticity, pooled for all networks. (B) The drivers (columns of *S*), which quantify the modulations of control ensembles (responsiveness, pliancy, choice) that capture each PC (pooled for all network classes). (C): The variance-weighted drivers for the three control ensembles, computed separately for the three network classes (fast, intermediate and slow).

We next calculated the matrix *S* of weighting factors (*drivers*) for the firing rate components, describing what combination of adjustments to the control ensembles best accounts for the associated firing rate changes (Fig. 4B; for full description of this approach, see Methods subsection *Modulation of control ensembles by plasticity*). To interpret the drivers of control ensemble influence (Fig. 4B), it is important to note that positive (negative) coefficients correspond to changes in control ensemble activity in the same (opposite) direction as indicated by the loadings in Fig 3. The first driver corresponds to a large amplification of the responsiveness control ensemble, and hence a decrease in various forms of global inhibition in the CBGT network. The first driver is also associated with a boost to the choice control ensemble, increasing the bias towards the rewarded choice The second driver has a strong negative weight on the choice control ensemble and a positive weight on the pliancy control ensemble. The third, fourth and fifth drivers feature weaker effects, with small modulations of all three control ensembles. Based on this analysis across all of the networks, the overall modulation of the control ensembles due to plasticity, calculated as the weighted sum over all drivers, adjusted by the % of variance explained by each PC, is shown in Supp. Fig. S5B. All three control ensembles end up being boosted. This means that, to varying extents, the activity measures that comprise these ensembles change in the directions indicated by their loadings in Fig. 3. In this way the general trend is for the CBGT networks to become more responsive, yet less pliant, which together amount to an earlier onset of evidence accumulation without much change in boundary height. Coincident with this, we also see that the CBGT networks exhibit more of an emergent choice bias with learning.

Because of the difference in decision policies across the fast, intermediate, and slow networks, we recomputed the drivers separately for each network type. This was done by considering the firing rate differences (Δ*F* ) and calculating the *S* loadings for fast, intermediate, and slow networks separately (see Methods - section *Modulation of control ensembles by plasticity*). The explained variance for each of the three network types is shown in Supp. Fig. S5A, and their corresponding PCs and goodness of fits are shown in Supp. Fig. S6. As expected, the drivers show different changes across the network types (Fig. 4C). The driving factor corresponding to responsiveness is negative for fast networks, while remaining positive for the others. The pliancy and choice factors were positive for all three networks, but pliancy was by far the largest for fast networks and quite small for the other two network types. Referring to the DDM parameter changes associated with changes in control ensemble loadings (Fig. 3), we see that the decrease in loading of responsiveness and strong increase in loading of pliancy for fast networks would both promote an increase in boundary height, *a*. This aligns with the fact that, of the three network types, only fast networks show an increase in *a* over the course of learning (Fig. 2, Supp. Fig. S7). Overall, we see that the specific way that plasticity adjusts the weighting of the control ensembles to drive changes in decision policies depends on the initial tuning of the network. Since plasticity results from the sequence of decisions and rewards that occur during learning, we next investigate more directly how specific decision outcomes lead to this dependency.

### The influence of feedback sequences on driving of control ensembles

In the previous section we described the overall effects of cortico-striatal plasticity on control ensemble tuning. We now turn to analyzing the early temporal evolution of these effects by focusing on the initial two learning trials. Specifically, we examined the modulation of the control ensembles for different combinations of successes (i.e., rewarded trials; R) and failures (i.e., unrewarded trials; U) achieved by the first two consecutive choices. For this analysis, we implemented our usual DDM fitting process followed by CCA for networks that were frozen (i.e., with plasticity switched off) after two trials, and we grouped the results based on the sequence of choice outcomes. The drivers (combined columns of *S*) for each sequence of outcomes, U-U, U-R, R-U and R-R, are shown in Fig. 5A.

**Fig 5.**
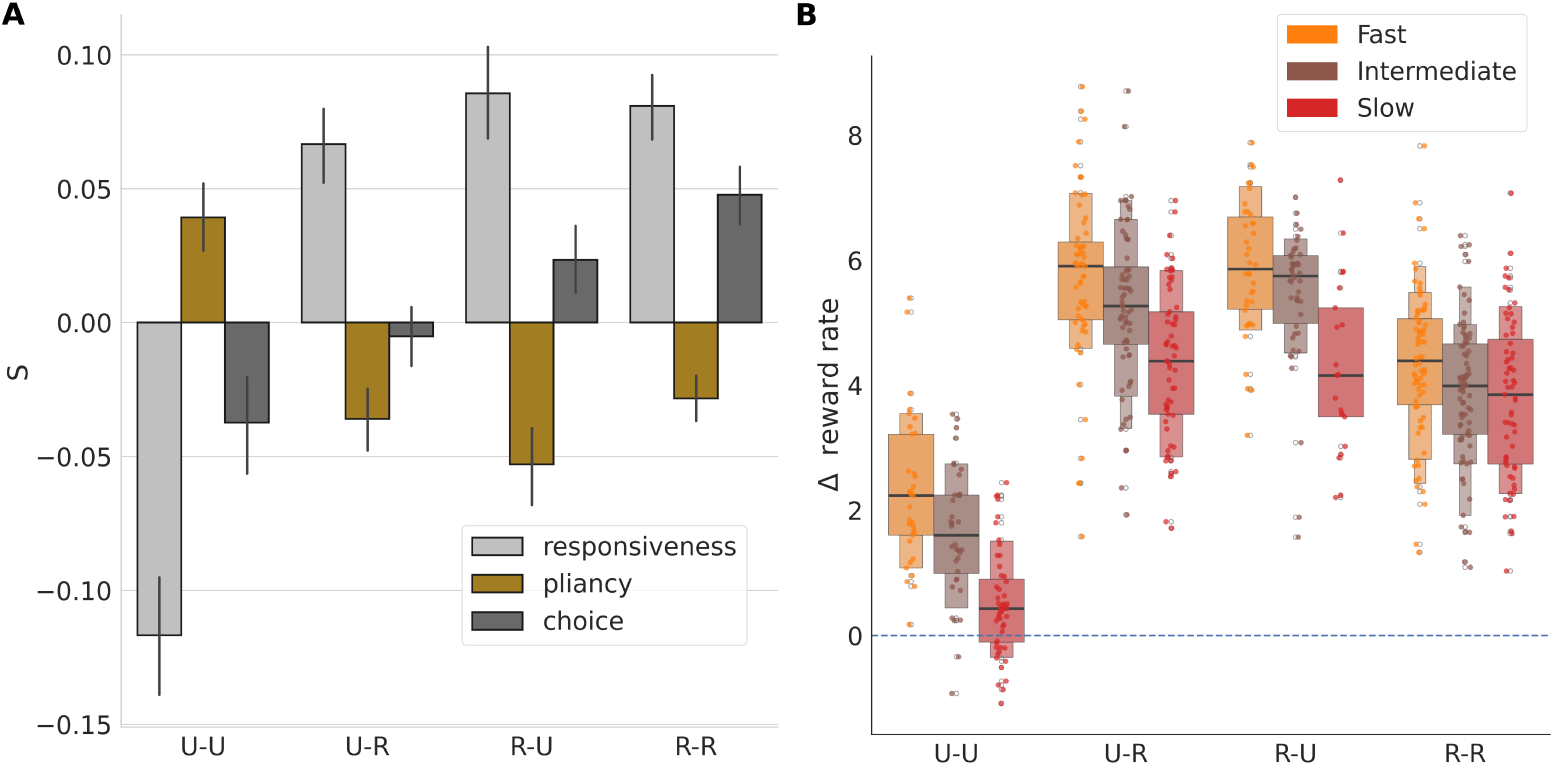
Suboptimal and optimal choices modulate control ensembles in opposite directions. (A) The modulation of control ensembles associated with various reward sequences encountered in two initial trials with cortico-striatal plasticity. U represents ”Unrewarded” and R represents ”Rewarded” trials. (B) The reward rate changes obtained by simulation of networks with synaptic weights frozen after various reward sequences occurred on two initial trials.

First, consider the case of networks that receive no rewards (U-U). Here we infer that the boundary height increases, due to a simultaneous decrease in driving of the responsiveness ensemble and increase in driving of the pliancy ensemble, both of which result in a boost of the boundary height. In addition, driving of the choice ensemble is reduced. Thus, two consecutive unsuccessful trials yield an overall increase in the degree of evidence needed to make a subsequent decision by simultaneously increasing the boundary height and decreasing the drift rate. Moreover, slow networks encounter U-U outcomes more often than other network classes in the first two trials (Supp. Table 1), which presumably constrains the increase in responsiveness and choice seen in these networks during learning (Fig. 4C). On average, however, fast networks make more mistakes than the other networks. This result, which we can display graphically in terms of the proportion of unrewarded trials, or mistakes, encountered after the first two plasticity trials (Fig. S7D), likely explains the negative loading for responsiveness and high positive loading for pliancy for fast networks shown in Fig. 4C.

In contrast, two consecutive successful trials (R-R, far right of Fig. 5A) produce largely the opposite effect. The influences of the responsiveness and choice ensembles increase, resulting in lower onset time and boundary height, along with an increase in the drift rate. This coincides with a weak change in pliancy. As a result, in the R-R case, the decision policy is tuned to include a decreased degree of evidence needed to make subsequent decisions.

Not surprisingly, the two mixed combinations of outcomes (U-R, R-U) have largely similar effects on the responsiveness and pliancy ensembles, regardless of the order of outcomes. In both cases the loading of responsiveness increases and that of pliancy decreases, resulting in less overall evidence needed to trigger a decision (by shrinking the boundary height, without much change in the onset time). However, when the first trial is unsuccessful (U-R) the influence of the choice ensemble decreases, while it increases when the first trial is successful (R-U). Indeed, looking at the progressive change in the choice ensemble across the four unique sequences of trials, it appears that early success (i.e., reward in the first trial) boosts the choice ensemble influence while early failure (i.e., unrewarded first trial) does the opposite. When these combined drivers are recomputed separately for each network class, the learning-induced modulations of the ensembles follow the same general trend (Supp. Fig. S8), with quantitative details depending on the network class.

The preceding analysis shows how the relative contributions of the control ensembles to the evidence accumulation process depend on trial outcomes. What are the results of these changes on the performance of the network? To illustrate these effects, we plot the distribution of changes in reward rates associated with each set of outcomes and separate by network types in Fig. 5B. Although all distributions are generally positive, there is significant variation in reward rate changes across the different feedback sequences (F(586, 3) = 254.4, p*<*0.0001). The reward rate also varies significantly with the network type (F(586, 2) = 46.8, p*<*0.0001), and the interaction term between network types and feedback sequences is significant as well (F(586, 6)=3.8, p = 0.001). Compared to all other conditions, the networks that made two consecutive unsuccessful choices (U-U) yielded the smallest changes in reward rates (values of all network types pooled together, all two-sample t(319) *<* -18.27, all p*<*0.0001). The two mixed feedback conditions (U-R, R-U) had higher growth in reward rates than the condition with two rewarded trials (R-R; all t(384) *>* 8.1, all p*<*0.001), because mixed conditions not only lead to strengthening of the correct choice but also weakening of the incorrect choice, unlike R-R which only leads to the former. In all cases, the trend was for faster networks to achieve greater increases in reward rate than slower networks. As expected, the impact of feedback sequences on reward rate is associated with underlying changes in both accuracy (Fig. S9A) and decision speed (Fig. S9B). Like reward rate, the increase in accuracy was highest for the mixed feedback conditions (U-R, R-U) due to the combined strengthening of the correct choice and weakening of the incorrect choice. Two consecutive unsuccessful choices (U-U) represents the only condition that leads to an increase in decision times, expressed as negative ΔRTs. This outcome is consistent with the increase in boundary height that occurs in this case, whereas all other feedback conditions lead to a decrease in decision times.

## Discussion

Adaptive behavior depends on flexible decision policies (*what* ), driven by CBGT networks (*where*) that shift their activity in order to maximize reward rate by coordinated adjustments of a set of underlying control ensembles (*how* ; Fig. 1). In this work, we focused on the *how* part of this process, employing a mapping upward in abstraction between a biologically realistic model of CBGT pathways and the DDM.

This approach helps to reveal the complex, low-dimensional structure of CBGT subnetworks that influence decision-making policies (Fig. 3). Specifically, we recapitulated recent results (30) showing the existence of three main CBGT control ensembles shaping decision-making that represent *responsiveness*, *pliancy*, and *choice* (direct vs. indirect pathway competition; Fig 3) and serve to regulate the process of converting evidence accumulation into action selection. We then showed how, within our model, driver mechanisms tune these control ensembles strategically during learning (Figs. 4 & 5) in order to maximize reward rate. Moreover, although they all optimize the same quantity (reward rate), we found that modulation of control ensembles differs across networks depending on their *a priori* decision policy (fast, intermediate, or slow). While plasticity increases responsiveness and choice in all networks, to varying extents, fast networks alone decrease responsiveness (Fig. 4C) and correspondingly increase boundary height (*a*; Fig. S7A). Put together, our results propose a new framework for understanding how subnetworks within CBGT circuits can dynamically regulate decision-making, driven by dopaminergic plasticity at the cortico-striatal synapses.

Perhaps the most surprising aspect of this theoretical analysis is the sophisticated adjustments that emerged from a simple plasticity mechanism acting on just one class of CBGT synapses. Dopaminergic learning at the cortico-striatal synapses was sufficient to push our naive networks from an exploratory decision policy to an exploitative policy that effectively managed the speed-accuracy trade off by maximizing average reward rate (Fig. 2). This behavior was recently observed in rats engaged in a perceptual learning task (32), indicating that reward rate maximization may be an innate behavior in many, if not all, mammals. However, the expression of this mechanism can vary depending on task contexts, such as differences in effort or feedback (15; 41). The rewards in our task that drove learning were based only on the accuracy of each selection. So how is it possible that rewards based only on accuracy can lead to an optimization of reward rate? The answer to this question lies in the architecture of the CBGT circuits. While synaptic plasticity in our model is limited to the cortico-striatal synapses, the resulting activity changes propagate throughout the entire CBGT network due to the synaptic coupling among the network’s interconnected populations. An emergent result from our simulations is that these cascading effects produce a subsequent reduction of decision times, even without any reward incentive that explicitly depends on speed. Thus, our model tends to act more slowly in the early phases of learning, but increases accuracy and speeds up decisions as learning progresses. This progression is similar to behavioral observations in rodents (32), non-human primates (42), and humans (43; 44). Our results predict that this complex behavior is a natural consequence of dopamine-dependent plasticity at the cortico-striatal synapses together with the architecture of the CBGT circuit.

Related predictions at the abstract level have been made by models that directly combine reinforcement learning with evolution in the DDM parameters (45; 46). These studies demonstrate that the drift rate depends on the difference in values between optimal and suboptimal actions, which increases with learning. In contrast, the boundary height is proportional to the effective values of the choices and typically shows a slight decrease as learning progresses. Another class of promising models are the reinforcement learning and racing diffusion (RL-RD) models, which can represent multi-choice decision-making with DDM-like accumulation. Some of these models (47) share a conceptual similarity with our CCA components in that they include q-values related to sum (Σ) and difference (Δ) elements. The RL-RD class of models offers alternative options for parameterizing learning data from our CBGT circuit model.

However, for the current work, we limited our analysis to estimating DDM parameters across learning stages to maintain consistency with our previous findings (30) and our current results on control ensembles in näıve CBGT networks.

Our primary goal with the analyses described in this paper was to decompose the circuit-level effects of plasticity that underlie adaptive reward rate maximization in terms of learning-related changes in the driving of the control ensembles. Based on the relation of the control ensemble loading to the evidence accumulation parameters (Fig. 3 & 4C), the effective learning-related changes result in shorter decision onset delays, higher rates of evidence accumulation, and variable changes in decision threshold as learning progresses (Fig. S6). On the shorter timescale of consecutive trials, each possible set of pairs of reward outcomes induces a specific adjustment of control ensembles in a way that increases subsequent accuracy and reward rate (Fig. 5, Fig.

S8). Interestingly, but perhaps not surprisingly, our results predict that mixed feedback, such as one rewarded and one unrewarded trial, will result in a stronger increase in reward rate than two consecutive rewarded trials. This finding is consistent with past results, as well as general intuition, on the benefits of exploration for effective learning (48; 49). It is, however, important to note that cortico-striatal plasticity may explain only a part of the decrease in decision speed seen in experiments. Additional boosts in speed may result from an agent’s increased confidence in the outcomes of its decision derived from other information sources (50). Moreover, an experimental paradigm that requires learning an explicit minimization of decision times may reveal other novel CBGT control ensembles, apart from those that we report here.

The existence of a small set of CBGT control ensembles, and the details of their components, represent some of the key predictions that emerge from our modeling study. Directly recovering these ensembles in real CBGT circuits would necessitate simultaneous *in vivo* recording of nine distinct cell populations across at least five distinct brain regions during a learning task. This is currently outside the scope of available empirical technology. While we hope that future experiments will test more focused aspects of our predictions, we can already extract relevant findings from the extant literature. For example, the predominant loadings in the responsiveness ensemble reflect the level of engagement primarily of input-level (cortical and FSI) components and inhibitory outputs (GPi) of the network, with higher loadings corresponding to less activation (Fig. 3). The increase in responsiveness associated with learning in intermediate and slow networks in our model therefore matches the suppression of activity in the subpopulation of striatal FSIs that was observed after learning in non-human primates (51). Interestingly, experiments have also found evidence for an earlier onset of activity in the striatum with the progression of learning in non-human primates (52). This is consistent with the decrease in onset time that arises via the learning-induced increase in the responsiveness and pliancy ensembles in all network classes in our simulations.

The pliancy ensemble, reflecting the influence of global striatal activity, including thalamic inputs to the striatum, as well as the influence of STN activity, is associated with the onset time and boundary height parameters; however, unlike the responsiveness ensemble, the pliancy ensemble has opposing loadings between onset time and boundary height. Thus, an increase in activity of the pliancy ensemble corresponds to an earlier onset of evidence accumulation, but with more information required to trigger a decision. This places an emphasis not on the collection of evidence itself, but instead on the agent’s willingness to be convinced by this evidence. It has been shown that an increase in the conflict between action values is associated with an increase in global STN activity (53; 54; 55), consistent with a strengthened driving of our pliancy ensemble. Also, because our simulations show an increase in efficacy of the pliancy ensemble with value-based learning (Fig. 4C) for fast and intermediate networks, we predict that the overall level of striatal SPN activity will increase as learning progresses. In contrast, activity in the GPi would decrease. The predominant contributions of this effect are predicted to occur in response to unrewarded trials (Fig. 5A). Consistent with these predictions, past studies have shown increases in striatal activity with learning (56). Related findings have been interpreted as being potentially linked to increased task attentiveness (57) or increased motivation (58; 59). Both effects are consistent with the lowering of onset time associated with our pliancy ensemble. Interestingly, increases in striatal activity, as measured via fMRI, have been found to be beneficial for learning in adolescents (60); our results suggest that such increases in the pliancy ensemble loading could relate to learning from mistakes (Fig. 5A, U-U case).

Finally, the choice ensemble, which corresponds to the degree of competition between direct and indirect pathways across action channels, is strongly associated with drift rate. Consistent with this relationship, single unit activity in dorsal striatum has been shown to reflect the rate of evidence accumulation and consequently preference for a specific response to a stimulus (61). At the macroscopic scale, we recently found that the competition between action representations in CBGT circuits, measured with fMRI, is indeed reflected in the drift rate in humans (7). At the causal level, a recent study with patients suffering from dystonia showed that deep brain stimulation in the GPi increased the likelihood of exploratory behavior, which was encoded as decrease in the drift rate (20). Whether deep brain stimulation (DBS) increases or decreases the output of its target area remains controversial (62; 63; 64). However, based on the loadings in the choice ensemble, we predict that the observed decrease in drift rate corresponds to increased similarity in activity across GPi neurons in different channels — a likely outcome if DBS similarly impacts all channels.

As learning occurs in our model CBGT network, the control ensemble loadings appear to co-evolve. The merit of the control ensemble idea is that it lets us decompose a complicated evolution process into interpretable components. Nonetheless, it can also be informative to consider combined effects that result from the simultaneity of changes in control ensemble loadings. As one example, we note that in non-human primates, stimulation of the caudate nucleus in the striatum reveals a negative correlation between drift rate and boundary height (65). Our model captures this early negative correlation in learning, where pairs of unrewarded trials decrease the loading of responsiveness and choice while increasing the loading of pliancy. This shift reflects overall striatal engagement across both channels, potentially mirroring the effects of stimulation, resulting in an increase in boundary height and a decrease in drift rate. In contrast, other outcome pairs produce the opposite effects. Over the longer course of learning in slow and intermediate networks, we observe an increase in drift rate and a decrease in boundary height, with responsiveness and choice ensembles playing a more prominent role. This trend suggests a gradual shift in importance from pliancy (e.g., overall striatal engagement) to responsiveness as learning progresses.

Overall, our results suggest how the low-dimensional substructure of CBGT circuits may adapt behavior during learning by adjusting specific aspects of the evidence accumulation process, thereby influencing the current state of a decision policy. Notably, we demonstrate that dopamine-dependent synaptic plasticity at cortico-striatal synapses, driven by choice-related reward signals, can strategically coordinate control ensemble activity to improve accuracy while reducing decision times, thereby maximizing reward rate. As we have discussed, these findings not only align with previous empirical observations but also offer clear predictions for future experimental investigations.

## Materials and Methods

### CBGT network

The CBGT model used in this work is a biologically constrained spiking network including neuronal populations from the striatum (dSPNs, iSPNs and FSIs), globus pallidus external segment (GPe), subthalamic nucleus (STN), globus pallidus internal segment (GPi), thalamus and cortex (excitatory and inhibitory components). For a two-choice task, each choice representation is implemented as a “channel” (24; 29; 30; 34; 66), so the model includes two populations of each neuron type except FSIs and inhibitory cortical neurons, which are shared across channels. On each trial, the two excitatory cortical populations receive excitatory synaptic inputs, representing evidence related to the available options, from a stochastic spike generator. This process has a baseline rate sampled from a normal distribution with a mean and standard deviation of 2.5 Hz and 0.06, respectively. To this baseline, we add a ramping component representing the presence of some stimulus or internal process that drives the consideration of possible choices. This component rises linearly until it reaches a maximum value (*f_target_* = 0.8), which was kept constant for all simulations in order to appropriately compare decision times. Specifically, we take

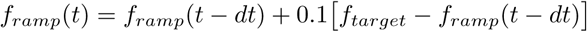

where *dt* is the integrator time step, such that the total frequency of inputs to the cortical populations evolves according to

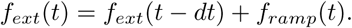

In all of our simulations, evidence for the two options, as represented by this frequency of inputs to the two cortical populations, was equally strong, such that changes in outcomes across conditions resulted entirely from learning downstream from the cortex. Specifically, the cortico-striatal projections to both dSPNs and iSPNs in the model were plastic and were modulated by a dopamine-dependent spike timing dependent plasticity rule (67; 68**?** ). On a trial, a choice was selected if the firing rate in the thalamic population within its action channel reached 30 Hz before the rate of the other thalamic population hit that level. The complete details of this network can be found in our methods paper (34).

### Characterization of networks before plasticity

In our previous work, we identified control ensembles based on extensive simulation of the CBGT network with each of 300 parameter sets selected using Latin hypercube sampling from among the ranges of synaptic weights that maintained biologically realistic firing rates across all populations (30). In that work, in which no learning occurred, however, the cortico-striatal projections to the choice representations (channels) were considered to be independent. Hence, some sampled network configurations were biased towards one of the choices. Because we studied the evolution of the control ensembles under plasticity in this work, we started with completely unbiased networks. Hence we resampled the networks from the joint synaptic weight distribution using genetic algorithms (see below) and isolated 300 networks that produced firing rates of all CBGT populations within experimentally observed ranges. The actual firing rate distributions are shown in Supp. Fig S1A. The networks before plasticity showed a diversity of reaction times (RTs, Supp. Fig S1B). The RT distribution was divided into 3 equal tertiles and used to define ”fast” (orange), ”intermediate” (brown) and ”slow” (red) networks. All of the networks before plasticity showed chance levels of accuracy (Supp Fig S1C).

### Genetic algorithms

The DEAP library (69) was used to run a genetic algorithm (GA) designed to sample CBGT networks with parameters from the ranges used in our previous work (30). Two additional criteria were used for the optimization function of the GA, namely (a) the network should produce trial timeouts (when no action was selected within 1000 ms) on fewer than 1% of trials, and (b) the network should be cortico-basal-ganglia driven; that is, the correlation between cortical activity and striatal activity should be positive. The first criterion ensured that we had ample decision trials included in the data, as needed to appropriately fit the DDM parameters (timeouts are dropped before fitting the DDM parameters). The second criterion ensured that the networks did not operate in a cortico-thalamic driven regime, in which cortical inputs alone directly pushed thalamic firing over the decision threshold.

The range for each parameter specified in past work (30) was divided into 30 bins and this grid was sampled to create populations. The indices of each bin served as a pointer to the actual values of the parameters in the ranges considered. The GA uses these indices to create, mate and mutate the populations. This ensures that the values of parameters remain within their specified ranges. For example, suppose that parameter *A* has range (-2.0,2.0) and parameter *B* has range (-0.3,1.0) and these ranges are each divided into 5 bins. The grids for parameters *A* and *B* will be:

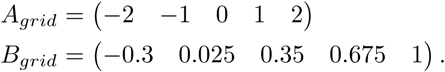

If individual population members have indices *ind*_1_ = (0 1) and *ind*_2_ = (4 0) for (*A, B*), then they have (*A, B*) = (−2, 0.025) and (*A, B*) = (2, −0.3), respectively. Supposed that the individuals mate by crossing over the 1st and 2nd elements. Then *ind*_3_ = (4 1) with parameter values (2, 0.025) and *ind*_4_ = (0 0) with parameter values (−2, −0.3). The individuals *ind*_3_ and *ind*_4_ are included in the next iteration of evolution.

New individuals created from mating were used to overwrite the original individuals that were mated together (*cxSimulatedBinary* ). The individuals could also mutate by shuffling of the indices of the attributes (*mutShuffleIndexes*) with a probability of 0.2. After a round of mating and mutation, tuples of two values for each individual, namely the % of timeouts and the Pearson’s correlation coefficient between cortical and striatal activity, were compared to select the individuals for the next round of evolution. The selection algorithm that was used was tournament selection (*selTournament* ) of size 3, which picked the best individual among 3 randomly chosen individuals, 10 times, where 10 is the size of the population of networks in every iteration of the GA. During every iteration, any network configuration that met the criteria (a) and (b) above was saved as a correct solution. The GA was run for 2000 iterations or until 300 solutions were found, whichever was sooner. Post hoc, we confirmed that the firing rates of the members of the final, selected populations remained within the originally targeted ranges (Fig. S1).

### DDM fits

The DDM parameters were fit to each of the 300 selected networks independently using the HDDM package (35). In order to ensure that the HDDM fits describe the choice and RT distributions well, we compared the post-predictive distributions of the network simulations with those generated by the corresponding DDM parameters, both before (Fig. S2A) and after (Fig. S2C) plasticity. The quantile-quantile plots for percentiles 5, 10, *. . .*,90, 95 show a significant and very high correlation between the network-generated and DDM-generated data (Fig. S2B,D).

### Accuracy, RT and reward rate manifolds

The manifolds shown in Figure 2A-C were generated by simulating the DDM with all combinations of drift rate (*v*) and boundary height (*a*) values that the naive CBGT networks can show before and after plasticity. The values of RTs, accuracy and reward rates were averaged over 15 seeds of 200 trials each.

### Upward mapping

The DDM parameters and activity of the CBGT nuclei for our 300 network configurations, before plasticity, were used to identify CBGT control ensembles through canonical correlation analysis (CCA), as was also done in our previous work (30) and is illustrated in Fig 3. The CCA scores were calculated using *k*-fold validation (*k*=4), where the 300 networks were divided into groups of 4 (75 networks each) and a CCA score was calculated for each of the groups. The CCA scores for actual data were compared with a shuffled version of data (firing rates and DDM components for 300 networks) and the set of components giving rise to the maximum CCA score, which we found to include three elements as in our previous work (30), were selected.

### Modulation of control ensembles by plasticity

We used a single approach to compute a set of effective drivers of the control ensembles either from the full collection of CBGT networks or from one of the network subtypes (fast, intermediate, or slow) that we considered. Let *X* ∈ {all, fast, intermediate, slow} denote the class of networks being used. From the set of vectors of changes in CBGT firing rates computed by subtracting firing rates before plasticity from those after plasticity (Δ*F_X_* ), we extracted 5 principal components (PCs) that together explain at least about 90% of the variance (Fig. 4A, Supp. Fig. S5A). Δ*F_X_* was then projected onto these 5 PCs to form the target matrix *P_X_* . Specifically, we computed

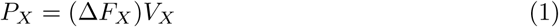

where the 5 PCs comprise the columns of *V_X_* . Note that *P_X_*is an *n* by 5 matrix, where *n* is the number of firing rate data vectors used. Δ*F_X_* was also projected onto the three control ensemble components obtained from the full collection of baseline networks before plasticity, via the mapping

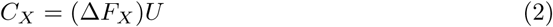

where the components of the 3 control ensembles form the columns of *U* , such that *C_X_*is an *n* by 3 matrix. Finally, we found the least squares solution *S_X_* , representing the element in the range of *C_X_* that is closest to *P_X_* , from the normal equation

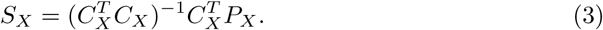

The least squares solution *S_X_*is a 3 × 5 matrix independent of *n*. The columns of *S*_all_ are displayed in Fig. 4B. The sums of the columns of the appropriate *S_X_* , each weighted by the percent of variance explained, comprise Figs. 5C and S7 (*X* = fast, *X* = intermediate, and *X* = slow), as well as Figs. 5A and S5B (*X* = all).

### Reward rates

The reward rate was calculated as:

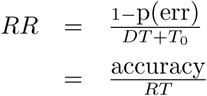

where p(err) denotes the error rate and where the reaction time, *RT* , is the sum of the decision time, *DT* , and the additional non-decision time that arises within each trial, *T*_0_, which in our analysis is ascribed to the onset delay represented by the DDM parameter *t*.

### Plasticity stages

The effect of plasticity on the network was studied at four stages: (a) after 2 trials of plasticity, (b) after 2 additional trials (total 4) of plasticity, (c) after 2 more additional trials (total 6) of plasticity, (d) after 9 additional trials (total 15) of plasticity. The state of the network was frozen at each of these stages by suspending the plasticity, so that we could use the frozen network to perform probe trials. The choices and reaction times from the probe trials were used to calculate DDM parameters and reward rate distributions for each stage of plasticity, based on upward mapping and CCA, and thus to generate the trajectories in Fig. 2, the time courses in Fig. S7, and the 2-trial results in Figs. 5, S8, and S9.

### Data sharing

The network codebase utilized in this study can be found on our GitHub repository and accessed at https://github.com/CoAxLab/CBGTPy/blob/main. Detailed installation instructions and a comprehensive list of implemented functions can be found in the README.txt file within the repository. All datasets generated and analyzed during the course of this research, along with a demonstration demo will be openly available on GitHub at https://github.com/jyotikab/CBGT_maximize_RR.

We thank all members of the exploratory intelligence group for their helpful comments on the manuscript. JB is supported by ANR-CPJ-2024DRI00039. TV, JB and JER are partly supported by NIH awards R01DA053014 and R01DA059993 as part of the CRCNS program. JER is partly supported by NIH award R01NS125814, also part of the CRCNS program.

---

**Fig S1.**
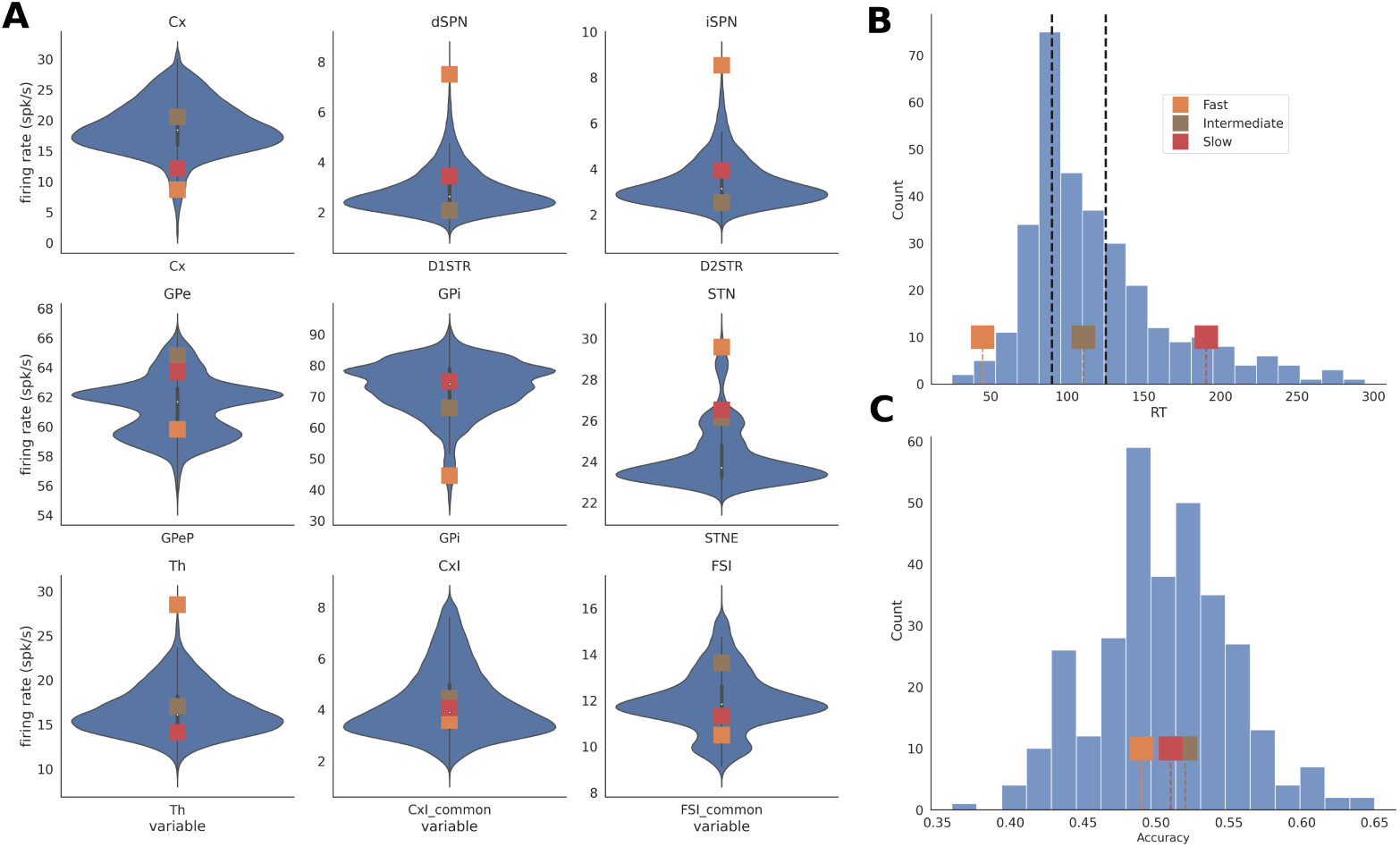
Network firing rates, RTs, and accuracy before plasticity. A: The distributions of average firing rates for the 9 CBGT regions based on 300 networks. An average was calculated for each population over the whole simulation time. One example each from three categories of network – fast (orange), intermediate (brown) and slow (red) – are marked on the distribution. B: The networks before plasticity were categorized as fast, intermediate and slow based on a tertiary split of the reaction time (RT) distribution (vertical dashed linebs). The RTs for the exemplar fast (orange), intermediate (brown) and slow (red) networks are marked. C: The average accuracies of all 300 networks. The accuracy distribution is centered around 50% (0.5) because the networks had not yet undergone plasticity.

**Fig S2.**
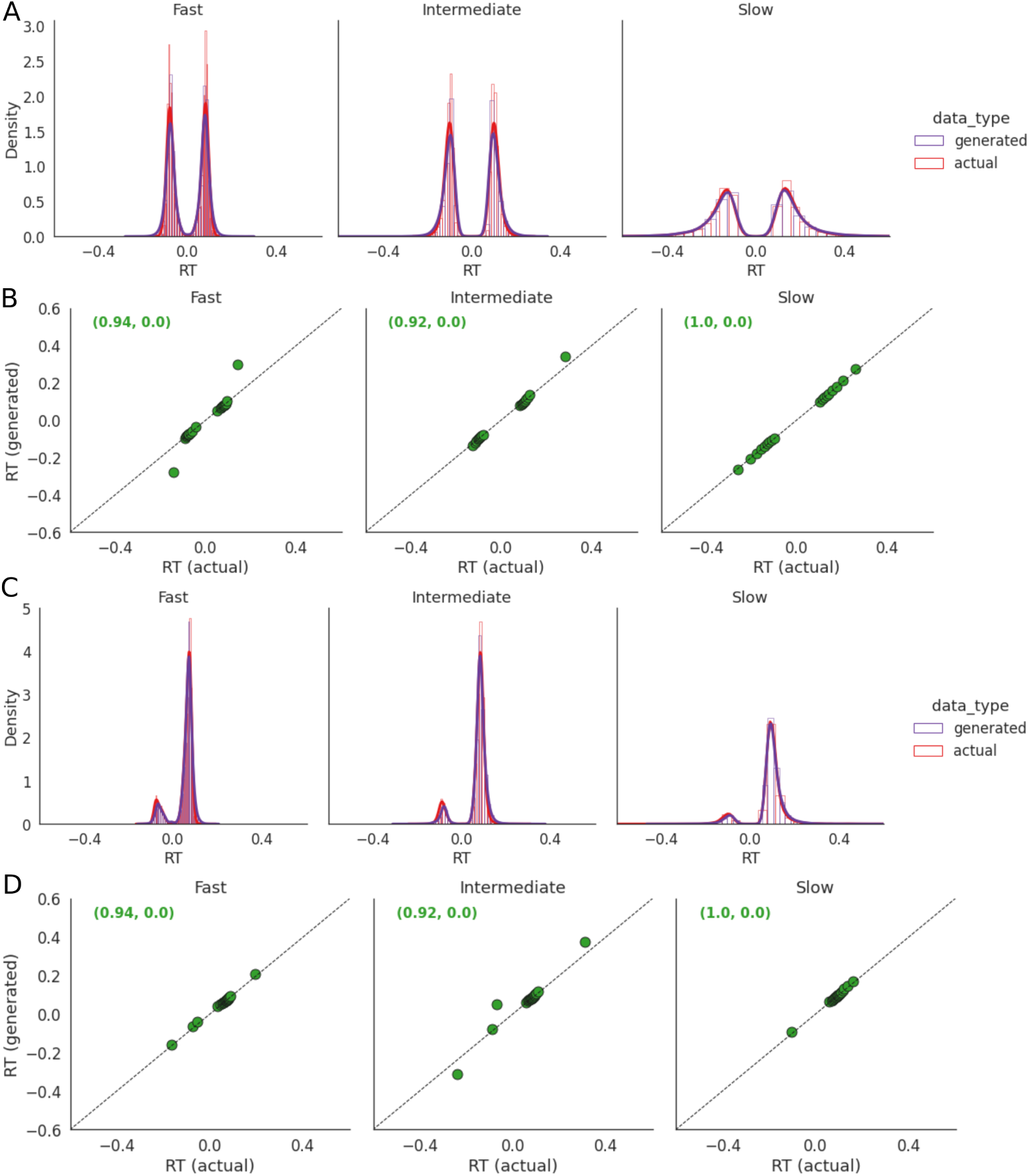
DDM fits for 300 networks before and after plasticity. A: The post-predictive choice (i.e., split between positive and negative RTs) and RT distributions from the naive (before plasticity) network simulations (red, “actual”) and distributions generated by the DDM parameters fitted to the data (purple, “generated”) separately for fast, intermediate and slow networks. Note the near-symmetry of the two RT peaks for the two choices (left → positive, right → negative). B: Quantile-Quantile plots for the distributions shown in A for percentiles in steps of 5 (i.e., 5, 10…90, 95). The Pearson correlation and p-value between the actual and generated data are annotated in green. The Pearson correlation was significant for all three network types (0.94, 0.92 and 1.0 for fast, intermediate and slow networks, respectively). C: Same as A but after plasticity with the left choice (positive RTs) rewarded. D: Same as B but after plasticity.

**Fig S3.**
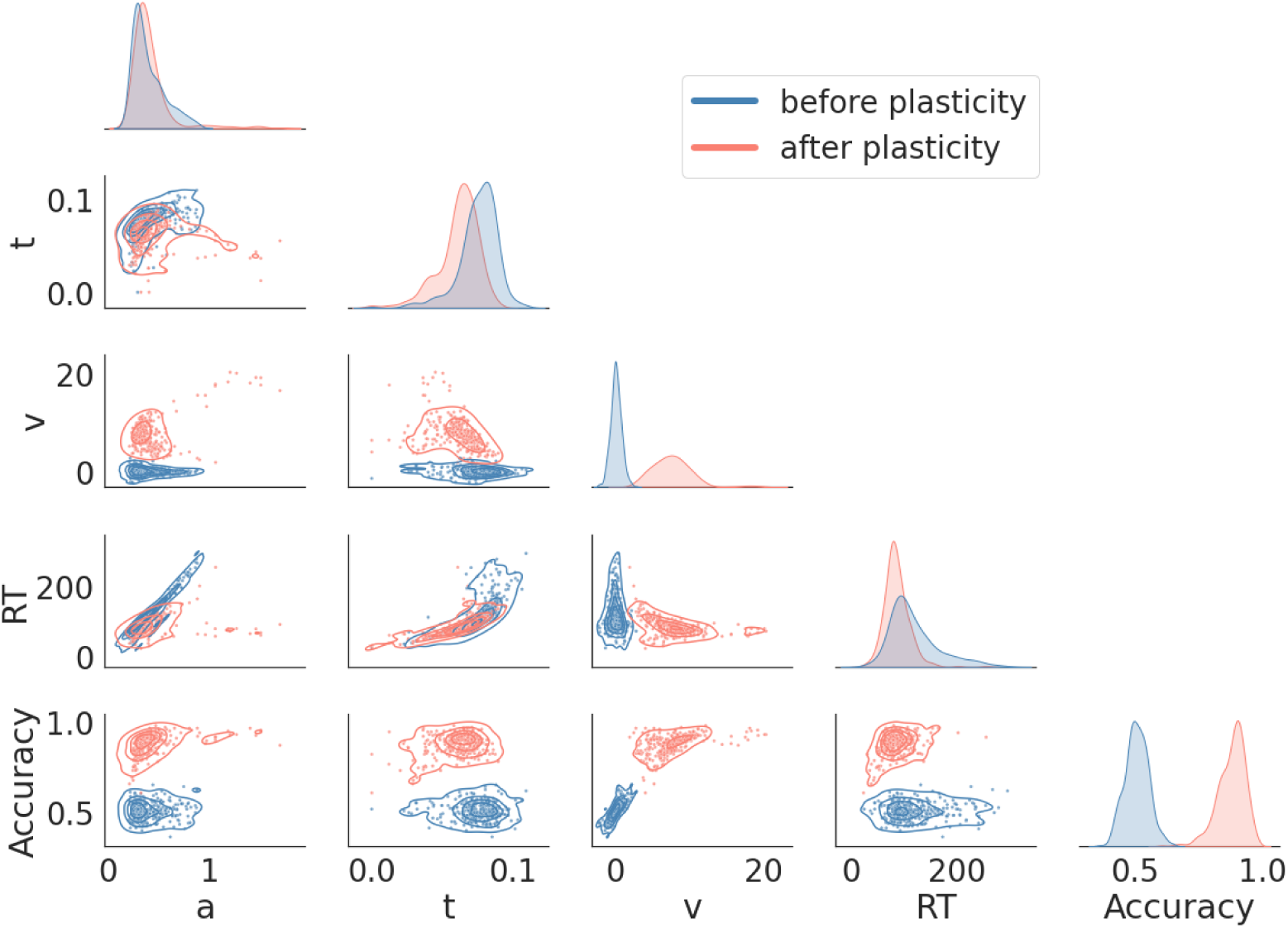
Comparison of DDM and behavioral measures for all 300 networks before (blue) and after (pink) plasticity. The subplots on the diagonal represent the marginal distributions for DDM parameters (*a*, *t*, *v*) and behavioral features (RT and accuracy). The onset delay (*t*) shows a decrease, the drift rate (*v*) shows an increase, RTs show a decrease, and accuracy shows an increase after plasticity. The off-diagonal subplots show the pairwise covariances.

**Fig S4.**
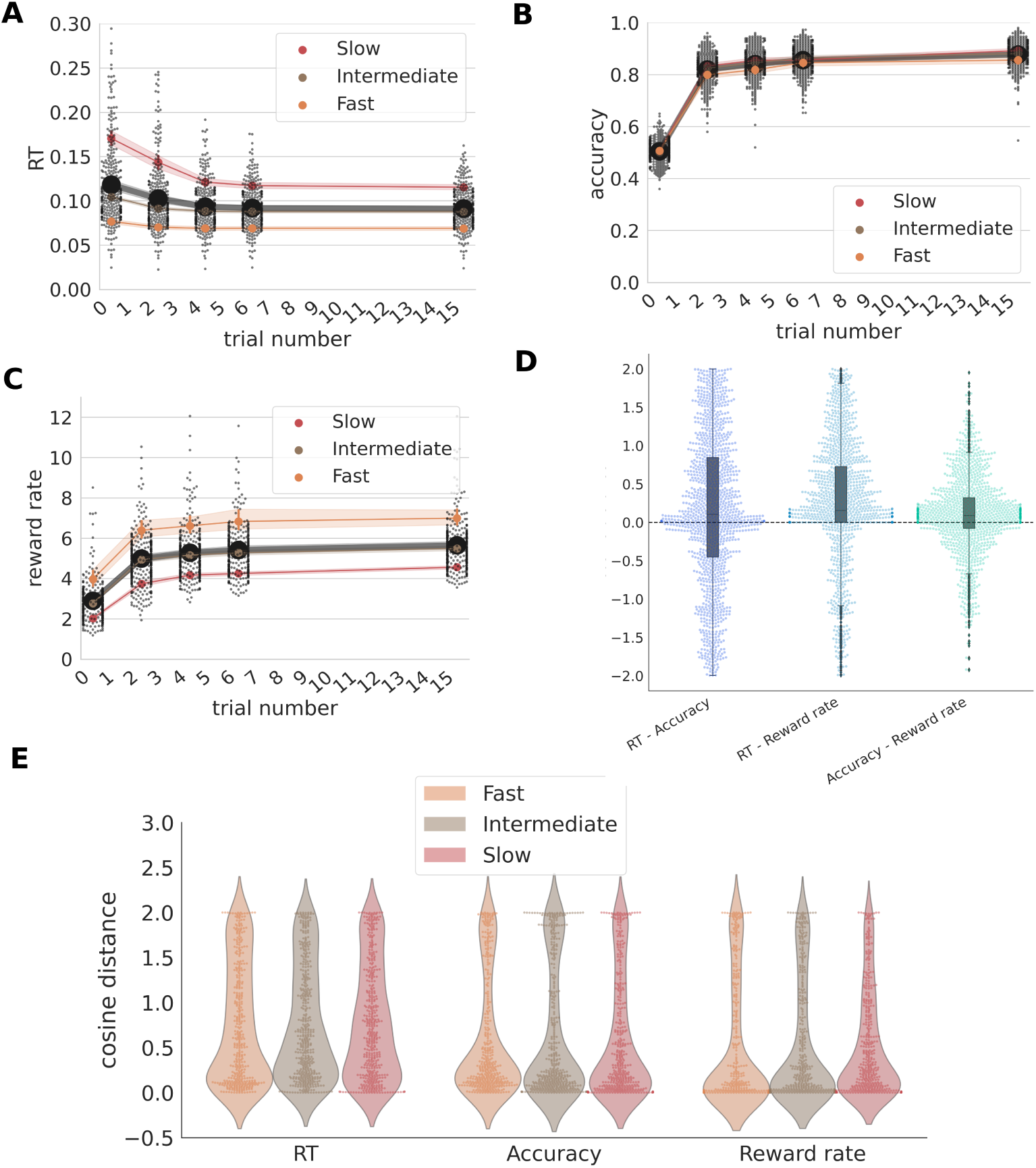
Evolution of behavioral measures for 300 networks over 16 trials with plasticity. A: Network behavior was assessed after each of 2, 4, 6, 9 and 15 trials. The RTs steadily decreased for all three network categories: fast (orange), intermediate (brown) and slow (red). The average over all 300 networks also showed a steady decrease as shown in black markers and lines. B: The accuracy for the three categories of the networks and the average over all 300 networks increased with plasticity. C: The reward rate for three categories of network and the average over 300 networks increased with plasticity. D: The distribution of differences in cosine distance, measured relative to the direction of greatest increase, for changes in RT vs accuracy, RT vs reward rate, and accuracy vs reward rate for all 300 networks and all stages of plasticity. The comparisons with reward rate yield distributions skewed to significantly above 0, suggesting that the cosine distances are lowest for reward rates. E: Absolute cosine distance distributions shown separately for the three network classes, fast (orange), intermediate (brown) and slow (red).

**Fig S5.**
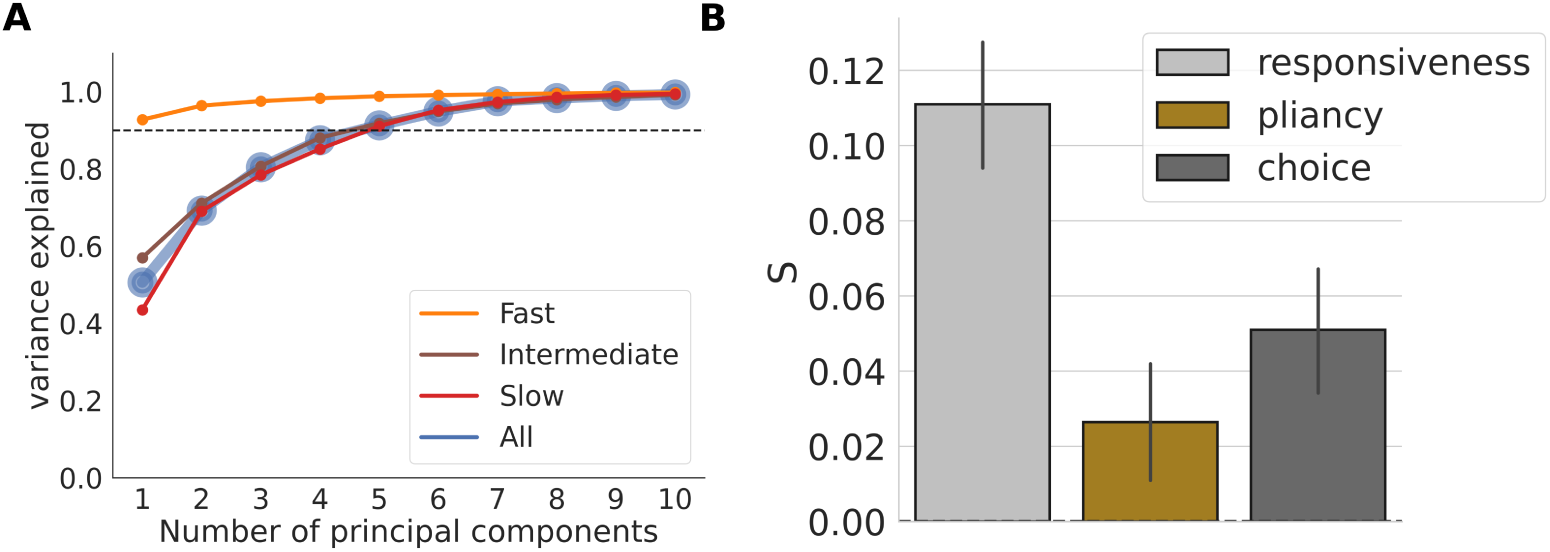
The least squares solution. *S* **pooled over the network types.** A:) Cumulative variance explained by the first 10 principal components (PC) derived from the changes in firing rates from before to after plasticity. The dashed line indicates 90% of variance explained. The analysis was done for all the networks pooled together (blue line) and separately for fast (orange), intermediate (brown) and slow (red) networks. For all networks pooled together as well as the separated slow and intermediate networks, the first 5 PCs explain more than 90% of the variance, whereas for fast networks 1 PC suffices. B: The weighted sum of the columns of *S* (see main text, Fig. 4B), pooled over all three network classes (fast, intermediate and slow), shows that the observed changes in firing rates correspond to increased loadings of the responsiveness, pliancy and choice ensembles of the CBGT network, to differing extents.

**Fig S6.**
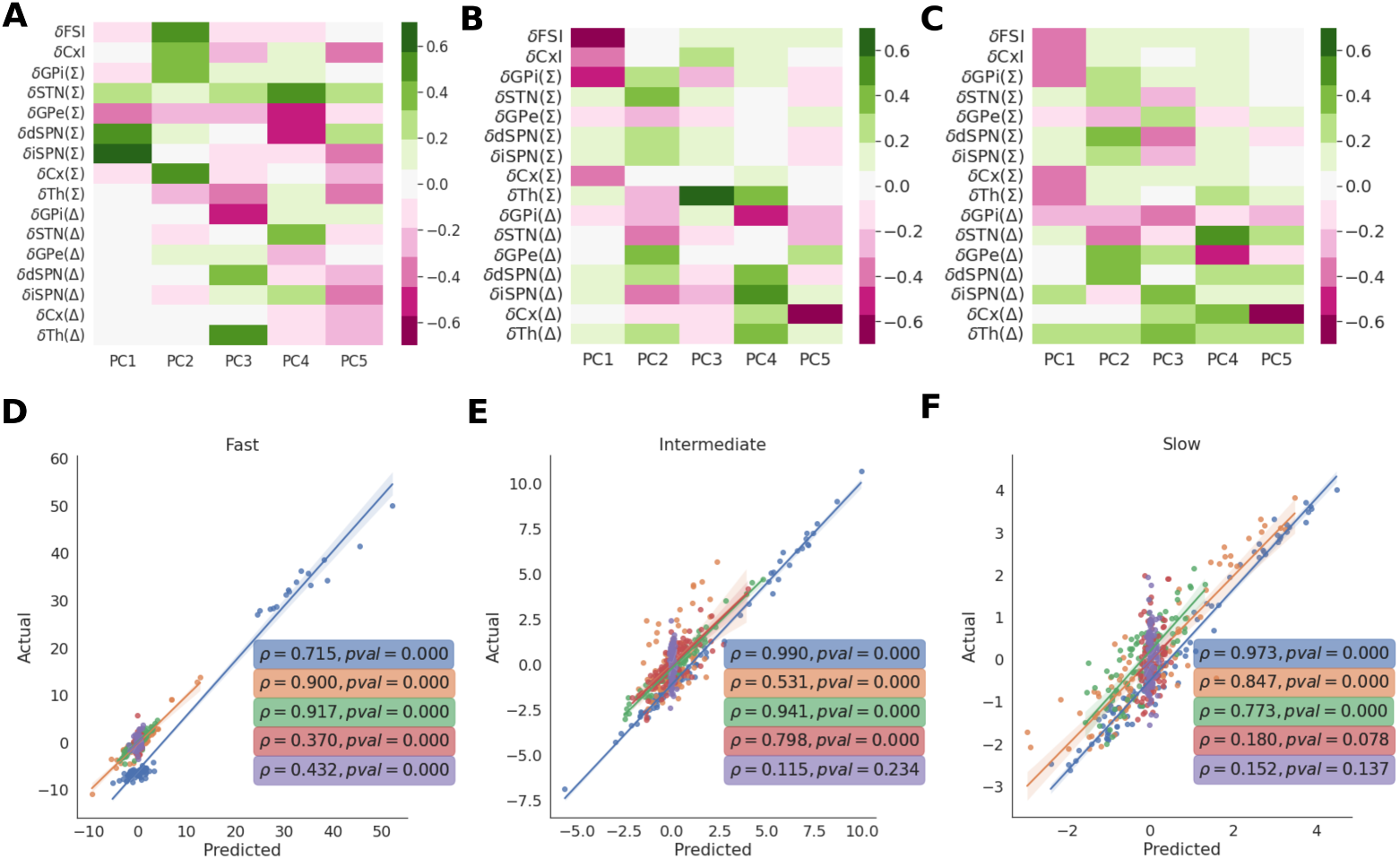
Reconstruction of firing rate changes from the least squares solution. *S* **for the three network classes.** (A) The first 5 PCs for the firing rate changes in the fast networks. Although the 1st PC explains around 90% of the variance for fast networks, we used 5 PCs to calculate *S* coefficients (Fig 4C) to be consistent with slow and intermediate networks (Supp. Fig. S5A). (B,C): Same as (A) for intermediate and slow networks, respectively. **(D-F)** The dot products of the CCA component vector (*C*) with each of the 5 columns of *S*, the least squares solution of *P* = *CS*, provide an approximate reconstruction of the 5 PCs of the changes in firing rate from before to after plasticity, (Δ*F* ). The quality of the reconstruction was checked by projecting Δ*F* onto the original PCs for each network (marked as *Actual* on y-axis) and comparing the results with the projections of Δ*F* onto the reconstructed PCs (marked as *Predicted* on x-axis). The goodness of fit is calculated as the Spearman rank correlation (*ρ*) between the actual and predicted values. For fast networks (D), the rank correlations (*ρ*) are high and significant (*p <* 0.0001) for all of the PCs as shown, suggesting that the reconstruction is excellent. For intermediate networks (E), the rank correlations are significant for all PCs except the 5th PC. For slow networks (E), the rank correlations are significant for all except 4th and 5th PCs.

**Fig S7.**
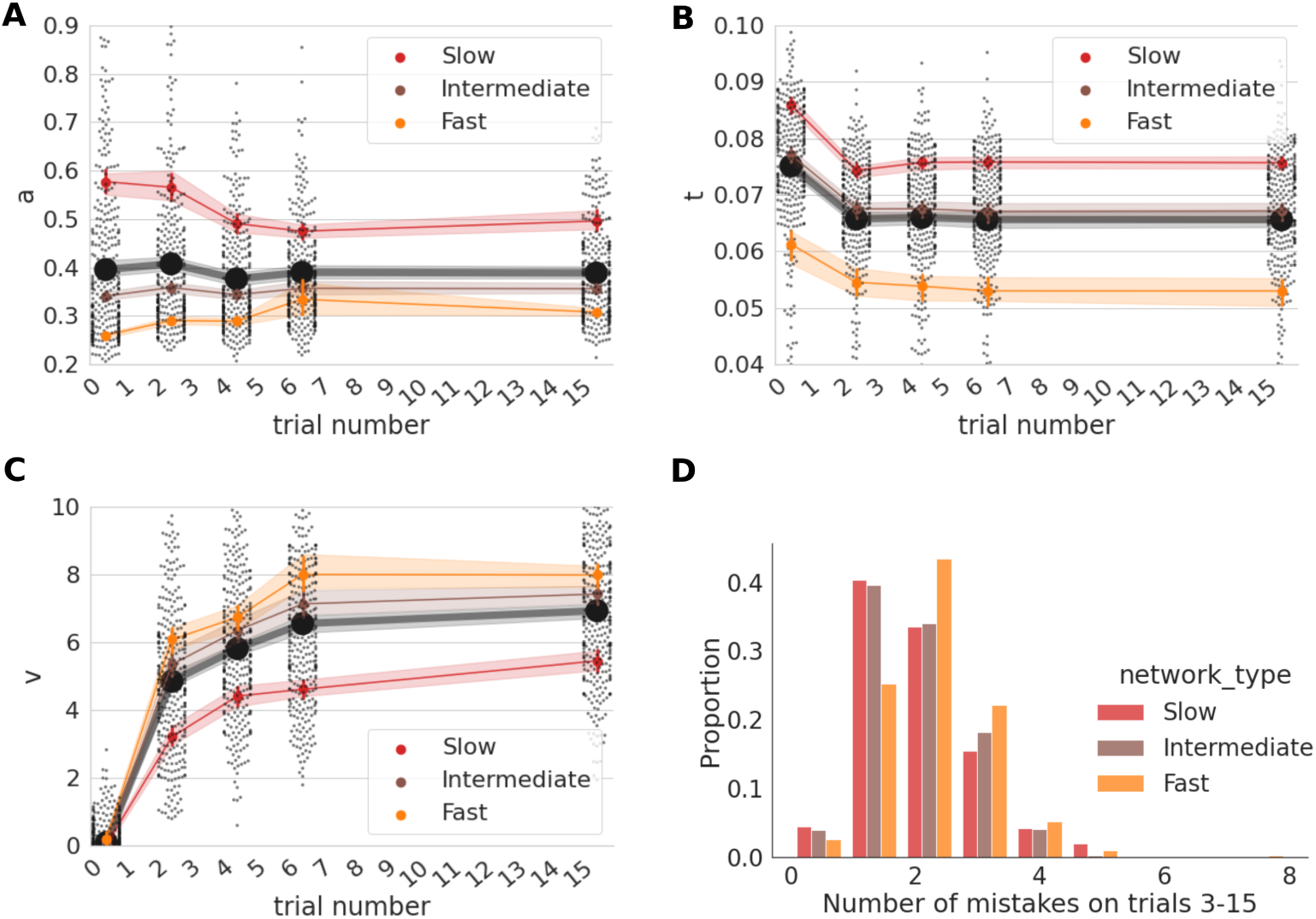
Evolution of DDM parameters with plasticity. (A) The change in boundary height (*a*) due to plasticity is dependent on network type: slow networks (red) show a decrease, intermediate (brown) show little change, and fast (orange) networks show a slight increase. The mean over all networks is shown by large black circles. (B) All network types show a decrease in decision onset time (*t*) due to plasticity. (C) All network types show a strong increase in drift rate (*v*) due to plasticity. (D) Fast networks make more mistakes on average. The histograms show the proportion of unrewarded (“U”) trials encountered by all the three network classes after the first two plasticity trials.

**Fig S8.**
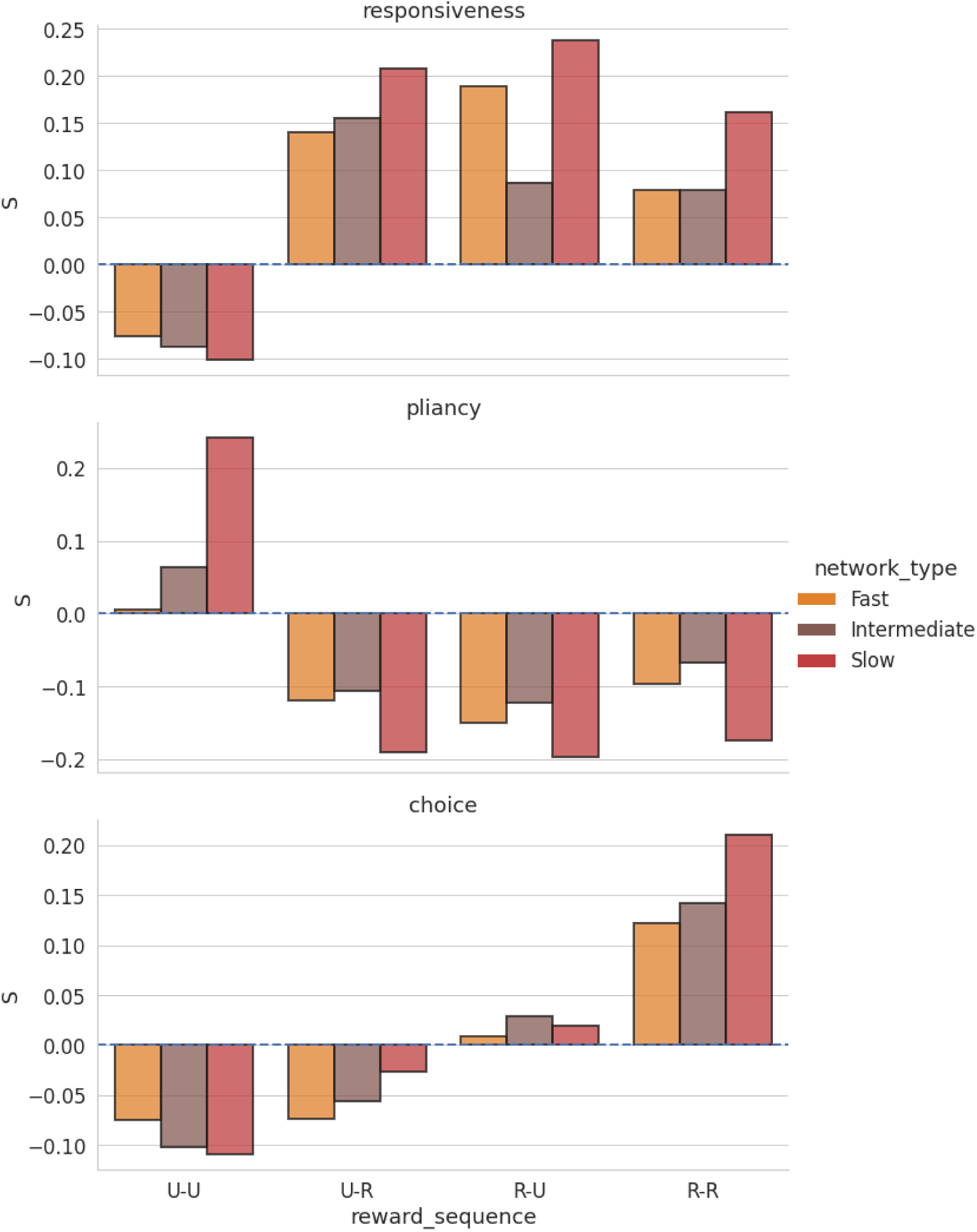
Effect of reward sequences on the weighting coefficients. *S* **for the three network classes.** The weighting coefficients *S* shown in Fig. 5A combine the three network types. The separated coefficients here show the same trends as the combined ones.

**Fig S9.**
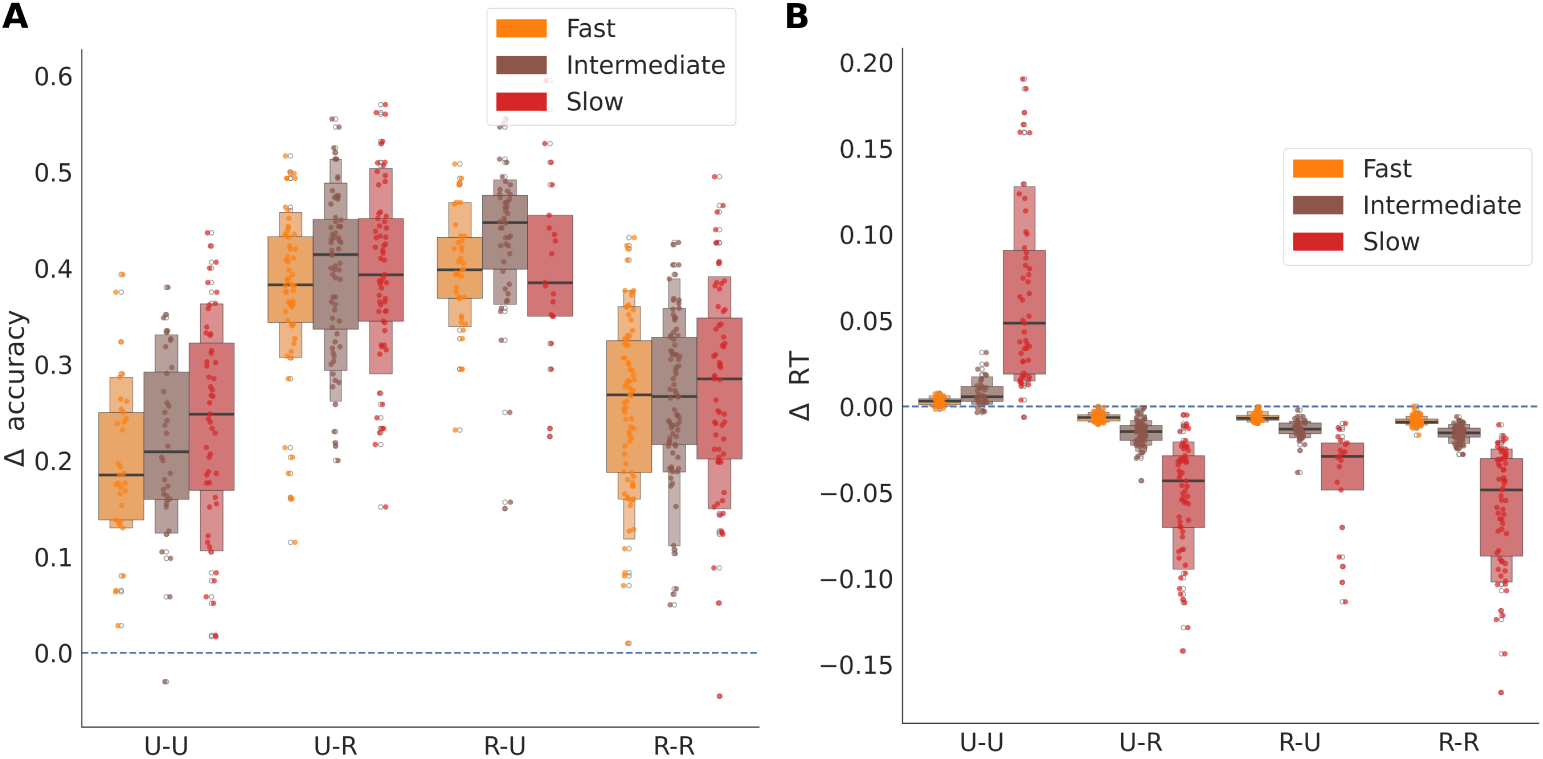
Effect of reward sequences on changes in accuracy and reaction times (RTs). (A) The change in accuracy showed an increase in all cases, but to different extents. The highest increase in accuracy was for one rewarded and one unrewarded trial (U-R and R-U), due to strengthening of the cortico-striatal projection to dSPNs of the optimal choice along with strengthening of cortico-striatal projections to iSPNs of the sub-optimal choice. (B) The change in RTs after plasticity for the four outcome sequences. All sequences involving at least one rewarded trial yielded a decrease in RT, whereas the sequence with two consecutive unrewarded trials (U-U) induced an increase in RT.

**Table S1.** Percentage of first pairs of trials for which networks encounter each possible reward sequence. Slow networks encounter a higher proportion of two consecutively unrewarded choices (U-U) and fewer R-U sequences than intermediate and fast networks.

| Reward sequence \ Network type | Network type |  |  |
| --- | --- | --- | --- |
|  | Fast | Intermediate | Slow |
| R-R | 36.3% | 35.1% | 32.1% |
| R-U | 18.7% | 19.9% | 10.7% |
| U-R | 28.9% | 29.4% | 31.1% |
| U-U | 16.2% | 15.6% | 26.0% |

